# DiffPIE: Guiding Deep Generative Models to Explore Protein Conformations under External Interactions

**DOI:** 10.1101/2025.04.27.650875

**Authors:** Yanbin Wang, Ming Chen

## Abstract

In recent years, many foundation generative models have been developed to predict structures of molecules and materials. Although these foundation models have achieved great success, it is challenging to collect enough data to train foundation generative models. One such example is to predict protein conformations with protein-environmental interactions (PEI), such as interactions introduced by organic linkers or material surfaces. We propose a physics-guided route to extrapolate foundation models beyond their training domain. Our method couples a pretrained deep generative model with explicit, physics-based interaction potentials for PEI, steering sampling toward conformations consistent with external constraints without any retraining or fine-tuning. We demonstrate accurate and efficient conformation prediction of (i) cyclic peptide with organic linkers and (ii) peptide adsorbed on the gold surface. The generated structures serve as high-quality initial conditions for downstream simulations, providing a general, systematic approach to extend foundation models to proteins under system-specific environmental interactions.

## Introduction

In recent years, foundation machine learning (ML) models in the chemistry community have attracted much attention. These models aim to solve fundamental questions in chemistry with broad applications derived from the high generalizability of deep neural networks. ^1^ Many of these models have been developed to address challenging problems that include, but do not limit to, property prediction,^2–7^ structure generations,^8–16^ force field parameterization,^17–20^ and chemical reaction prediction. ^21,22^ One type of these foundation models, named generative models, aims to generate atomic structures of small molecules,^23–30^ biomolecules,^16,31–37^ and materials.^38–44^ Conventionally, structure generation has been a highly challenging task that can only be solved by optimization-based methods^45–54^ or physics-based simulation approaches, ^55–62^ while the development of the foundation generative models provides highly efficient solutions to the problem of structure generation. These developments significantly advance research fields such as drug development,^24,25,30^ protein structure prediction,^16,31–37^ and material designs. ^38–44^ However, it is questionable whether a foundation generative model is capable of efficiently exploring individual chemical spaces of small organic molecules, materials, biomolecules, and others. A simple estimation of the number of organic molecules made by C, H, O, N, S, and with a molecular weight less than 500, is about 10^60^.^63^ The number of possible Wyckoff sequences, a way to classify crystal structures, is approximately 10^18.64^ An estimate of proteins that possibly have been present since the origin of life is about 10^21^.^65^ The huge size of an individual chemical space already poses significant challenges in collecting high-quality data to train foundation generative models to explore the space. In fact, the success of protein foundation models is highly dependent on the accumulated high-quality protein structure data over decades.

It is even more challenging to generate the atomic structures of chemical systems in interactions with the environment. Using proteins as examples, the protein-environment interaction (PEI) includes possible covalent and / or noncovalent interactions with organic molecules,^66^ materials,^67,68^ soluble polymers,^69^ and membranes. The generation of protein-with-PEI conformations is crucial to understanding protein functions in a wide variety of applications in biosensing, antifouling, drug delivery, and drug design.^70–72^ Although highly important, methods for predicting protein-with-PEI conformations are limited. Although experimental methods such as X-ray crystallography,^73^ NMR,^74^ and cryo-EM^75^ have achieved great success in determining the atomistic structures of proteins, the direct application of these structural determination methods to protein with PEI is challenging. These limited examples of structural studies focus on protein-protein complexes,^76^ protein-nucleic acid complexes,^77,78^ protein-drug complexes,^79^ and cyclic peptides.^80^ However, resolving protein structures with many other PEI types, such as protein-material interactions, is a challenge. More often, protein conformations with these PEI are measured using ensemble-based methods such as IR, circular dichroism, and SAXS where obtaining atomic structures is still an open question.^81–83^ In addition to experimental approaches, theoretical modeling to resolve protein conformations is also challenging. Generic approaches to exploring proteins conformations under external interactions include all-atom (AA) molecular dynamic (MD) simulations and coarse-grained MD simulations. ^84–89^ All-atom MD simulations are often limited by simulation time scale and system length scale. Although CG models are accessible to larger system sizes and longer simulation timescales, improving the accuracy of CG models is still an open problem.^90,91^ The challenges of determining protein-with-PEI conformations become significant obstacles in collecting data to train a foundational generative model of proteins with PEI. Therefore, current foundation models can only predict protein structures with limited classes of PEIs such as protein-biomolecule complex or protein-drug complex.^92,93^ Thus, guiding existing deep learning models to explore protein conformations under external interactions represents a promising avenue for fully utilizing the potential of foundation models.

Recent studies have begun to explore similar directions,^94^ particularly through guided sampling using diffusion models.^95–97^ Diffusion models learn by progressively adding noise to data points and can generate new data through a reverse diffusion process.^98,99^ In particular, the reverse diffusion process, either its drift or its diffusion term, can be modified to manipulate the data points generated.^100,101^ Examples of such modifications include incorporating restraint potentials to ensure that all-atom structures generated are consistent with experimental measurements,^101^ enforcing the generation of structures containing specific motifs,^102^ and guaranteeing correct stereochemistry in protein–ligand complexes. ^25^ Some studies have also used AA force fields to refine protein structure generation.^95,97^ Building upon these ideas, we present a systematic framework, termed the “Diffusion Model for Proteins Interacting with the Environment (DiffPIE)”, to integrate PEIs into an existing diffusion model. The core idea is that a diffusion model implicitly learns a “statistical potential” for protein structures without PEIs. By adding explicit physical interactions between the protein and the surrounding chemical species to this statistical potential, we construct a new effective potential energy landscape. Incorporating this modified potential into the diffusion process allows the model to generate protein structures that account for external interactions.

We will demonstrate the success of the proposed framework with two representative examples: a peptide denatured on a gold surface and a peptide cyclized with an organic linker. In both cases, we show that PEIs can be effectively modeled using MD simulations of fragments of the protein, rather than simulating the entire protein with its environment. Furthermore, we demonstrate that incorporating physics-based potentials effectively guides the structure generation process. Short plain MD simulations are then used to relax the generated structures, illustrating that the DiffPIE model can identify high-quality initial structures for proteins with PEIs.

## Results

### DiffPIE: Guiding Protein Structure Generation with Protein-environment Interaction

Assume that a protein structure ensemble can be generated using a pre-trained foundation score-based diffusion model, which learns through a forward noise-adding process described by:

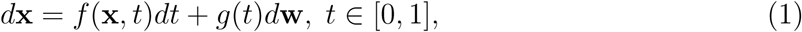

and generates new structures through a reverse denoising process:

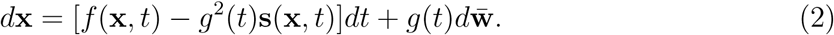

To simplify the discussion, we consider **x** in Eqs.(1) and (2) as representing atomic coordinates. As we will demonstrate later, DiffPIE can also operate with diffusion models that hybridize Cartesian coordinates and generalized coordinates. Here, *f* (**x**, *t*) and *g*(*t*) define the noise schedule, while **s**(**x**, *t*) is trained to approximate ∇**_x_** log *P_t_*(**x**), where *P_t_*(**x**) is the time-dependent marginal distribution in Eq.(2). The resulting generated structures follow a distribution *p*(**x**) learned from the training dataset, which aims to approximate the Boltzmann distribution of a protein without PEI, denoted as 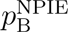(**x**). However, as illustrated in Fig. 1(c), *p*(**x**) can differ significantly from the true Boltzmann distribution of a protein under PEI, 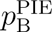(**x**). Consequently, it is challenging to study the thermodynamics of 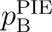(**x**) using structures generated solely from *p*(**x**).

**Figure 1:**
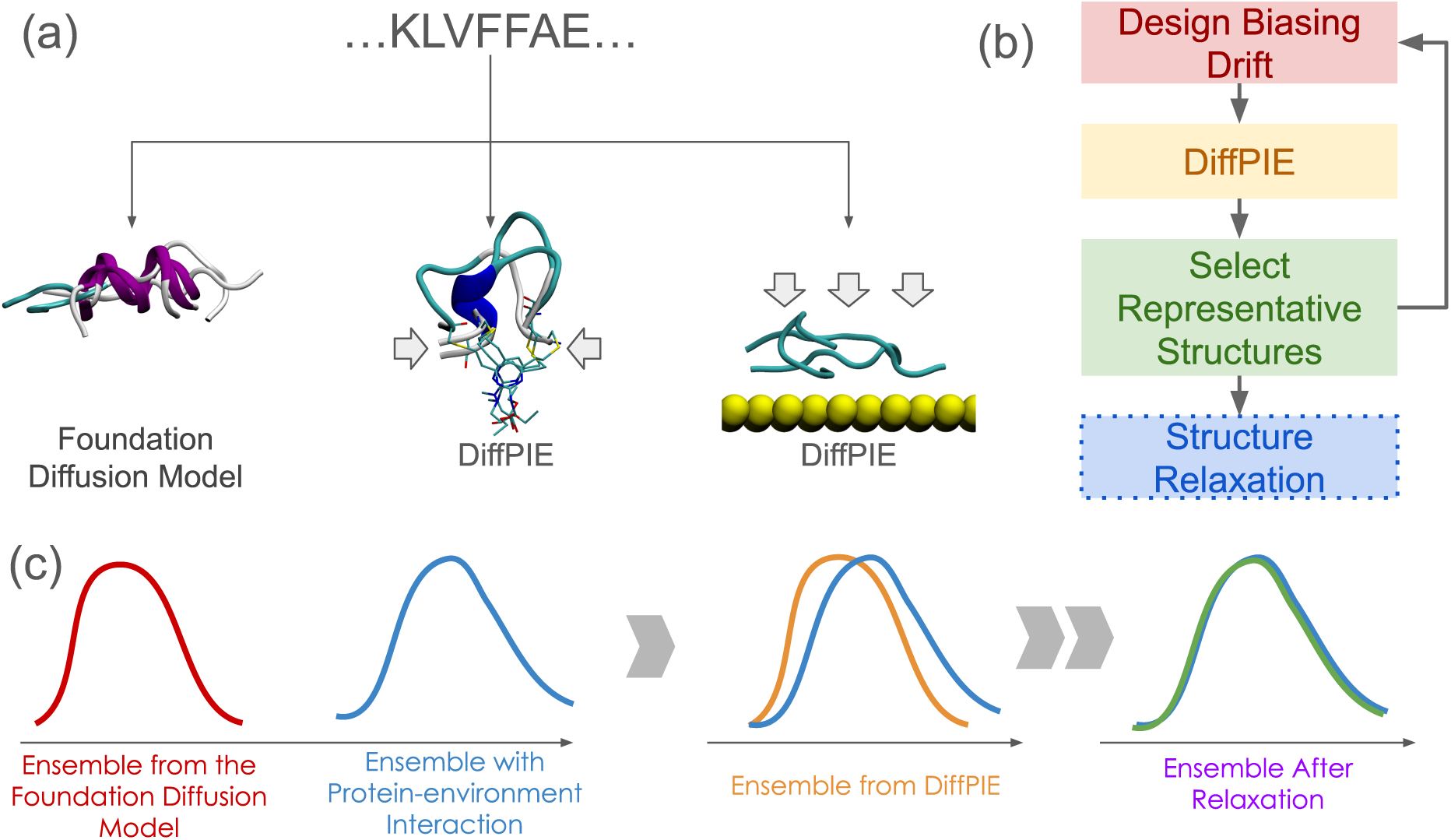
Panel (a) illustrates the concept of biasing protein structure generation using **F**_0_(**x**). The left bundle of structures was generated by the foundation diffusion model, Str2Str, with **F**_0_(**x**) = 0, corresponding to unbiased generation. The middle bundle shows structures generated with a **F**_0_ designed to mimic the restraint forces imposed by an organic linker. The right bundle presents structures generated with a **F**_0_ simulating adsorption forces, such as those from a surface interaction. Panel (b) summarizes the overall workflow of DiffPIE. Once a biasing force is designed, DiffPIE can be applied without any retraining. The distribution of generated structures can, in turn, inform adjustments to the hyperparameters in DiffPIE or refinements in the design of the biasing force. Although structure relaxation is not the primary focus of this paper, we demonstrate examples using short MD simulations for further refinement. Panel (c) illustrates how DiffPIE alters the ensemble distribution. The ensemble generated by the foundation diffusion model without bias (red) can significantly differ from the true protein ensemble under PEI (blue). By introducing the biasing force through DiffPIE, the generated ensemble distribution (orange) is shifted closer to the true ensemble. Additional MD relaxation can further improve the ensemble (green) to better approximate the true distribution.

To address this, DiffPIE generates a biased distribution *p*_bias_(**x**) that approximates 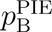(**x**), enabling rapid relaxation toward an ensemble representative of the true protein-environment system. DiffPIE achieves this by modifying the reverse diffusion process:

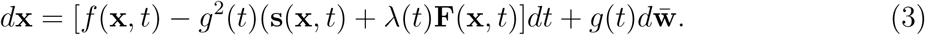

Here, **F**(**x**, *t*) is a biasing drift term that encodes protein-environment coupling, and *λ*(*t*), which changes with *t*, controls the strength y of the applied bias. A larger *λ* leads to stronger bias away from *p*(**x**). The details of designing *λ*(*t*) will be discussed in the “Method” section. In DiffPIE, **F**(**x**, *t*) is constructed by propagating a biasing force **F**_0_(**x**), modeled by fragmentation of the studied protein, back into the diffusion model, following an approach similar to the manifold constraint algorithms. ^103,104^ Users have flexibility to design **F**(**x**, *t*) suitable for a wide range of PEI, as illustrated in Fig. 1(a). Different choices of **F**_0_(**x**) allow the generation of protein structures consistent with different PEI scenarios. It is important to emphasize that DiffPIE does not require any retraining or fine-tuning of the foundation diffusion model. Instead, it only modifies the generation process by introducing the biasing drift **F**(**x**, *t*) during sampling. The DiffPIE framework is illustrated in Fig.1(b). We begin by designing a customized biasing force, with specific examples provided in following sections. In this study, we use Str2Str^16^ as the foundation diffusion model; however, in principle, any diffusion model can be incorporated. A key advantage of Str2Str is its flexibility in controlling the generated ensemble by selecting different seed structures and tuning hyperparameters. From the generated structures, we select representative conformations which can serve as new seeds or inform further hyperparameter adjustments. This iterative process is repeated until no additional new conformations are generated. Finally, the selected structures are subjected to relaxation through either structure minimization or short MD simulations, depending on the application.

### Sampling Cyclic Peptide Structures Containing Organic Linkers

P3-F is a cyclic peptide (Fig. 2(a)) that can interrupt interactions between the Nuclear Factor Erythroid-2-related factor (Nrf2) and the E3 ligase Kelch-like EH-associated protein 1 (Keap1) to counteract oxidative stress. ^105^ The sequence of P3-F is CDPETGECL. Two cysteines are cyclized by divinylpyrimidine with Michael addition between the vinyl group and cysteine (CYS) thiols.^105^ Here, we use P3-F as an example to demonstrate how to use

**Figure 2:**
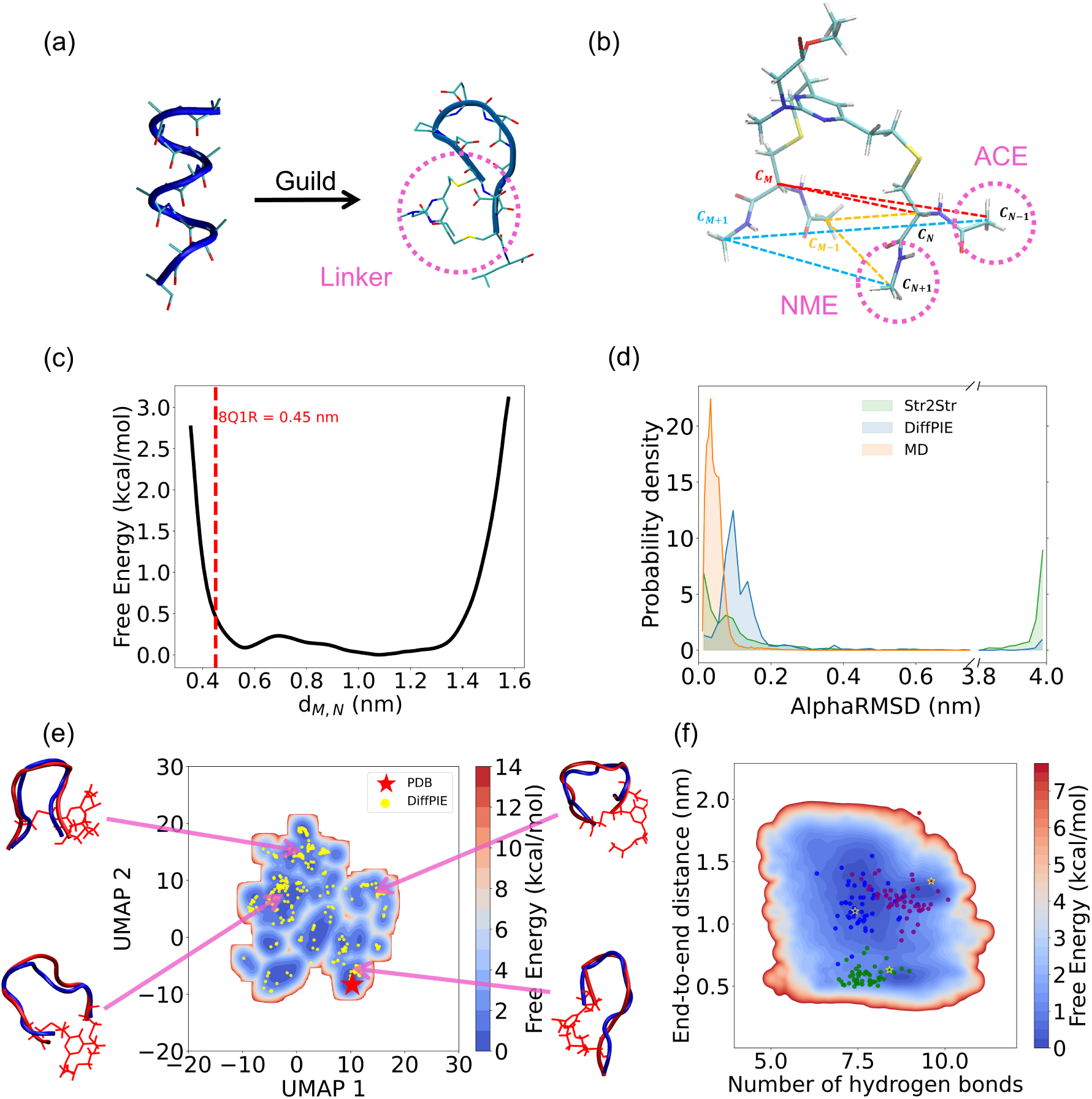
(a) DiffPIE is designed to guild the peptide (P3-F) sampling towards the structure with geometric restraint from linker molecule (PDB ID: 8Q1R). (b) Model molecule for **F**_0_: a staple with two CYS residues capped by acetyl (ACE) and N-methylamide (NME) to form dipeptide-like ends. Six dashed-line distances serve as collective variables (CVs), with mean forces yielding **F**_0_. (c) Free-energy surface (FES) of the C*α*–C*α* distance between the two CYS in the model, with the red dashed line from the PDB structure. (d) Effect of biasing force on sample distribution, shown as AlphaRMSD for Str2Str, DiffPIE, and metadynamics MD. AlphaRMSD measures the *α*-helix similarity of generated structures. Small AlphaRMSD means the structure is dissimilar to *α*-helix while large the structure is similar to *α*-helix. (f) Background FES from metadynamics configurations mapped via UMAP, with PDB (red star) and DiffPIE samples (yellow) projected onto the same space; representative structures from simulation (red) and DiffPIE (blue) are compared. (g) 10-ns unbiased MD relaxation from low-energy DiffPIE clusters, with initial structures (stars) and trajectories (dots) projected onto an FES of end-to-end distance and backbone hydrogen bonds.

DiffPIE to generate peptide structures constrained by organic linkers.Fig. 2(b) demonstrates how to construct **F**_0_. We first used MD to simulate a system of two cysteine dipeptides cross-linked by the linker molecule. The thermodynamic mean force of six distances (labeled with colored dashed lines in Fig. 2(b)) has been used as **F**_0_. Fig. 2(c) highlights a one-dimension free energy surface (FES) of the distance between two *α*-carbons (C*α*) in two cysteines. It is clear that the FES is highly anharmonic with multiple minima, suggesting that the conventional manifold constraint method, which uses a harmonic potential to apply restrictions to generated structures, is not applicable in this case. Fig.2(c) also highlights that the C*α*–C*α* distance in the PDB structure is not a minimum in the FES, suggesting that stable structures in this cyclic peptide should balance the internal energy of the peptide modeled by the foundation diffusion model and the bias energy of the linker modeled by **F**_0_. Fig.2(d) shows the impact of applying a bias drift. Structures generated by Str2Str without **F**_0_, DiffPIE, and a metadynamic^106^ simulation of the entire peptide are mapped in the space of AlphaRMSD.^107^ Str2Str generated a large number of helix-like structures, while these structures are almost not seen in the metadynamics simulation. DiffPIE significantly reduces the population of helix structures and highlights the conformation with a low AlphaRMSD, which is consistent with the conformations from metadynamics. We also examine the diversity of the generated structures in Fig. 2(e). The background FES of the Uniform Manifold Approximation and Projection (UMAP)^108^ CVs was generated after unbiasing the metadynamics simulation.^108^ We map the generated structures onto the same UMAP CVs as the scatter points. This clearly demonstrates that DiffPIE is capable of generating structures that visit most of the stable conformations. The PDB structure, denoted by the red point, corresponds to a metastable conformation. Since the PDB structure was resolved for the Keap1–P3-F complex, it indicates that the protein–peptide interaction is the main driving force stabilizing the P3-F PDB structure. Even though the PDB structure is not the most stable conformation in the metadynamics simulation, DiffPIE successfully generates similar structures, highlighting a possible application of DiffPIE in predicting cyclic peptide drug binding.^109^ Fig. 2(e) also shows that multiple structures generated by DiffPIE are comparable with stable structures sampled by metadynamics. In addition, we examine the quality of the generated structures and their captivity as initial structures for downstream sampling. First, stable conformations are included in the structural library generated by DiffPIE, as shown in Fig.2(f). Short MD relaxations initiated from the and purple structures stabilized within their respective conformations. In contrast, Fig. 2(f) shows a non-optimal green structure generated by DiffPIE that, although suboptimal, was able to rapidly relax to a similar conformation.

### Sampling Peptide Structures on Gold Surface

Gold nanoparticles have been widely used in biochemistry, with applications such as surface plasmon-enhanced spectroscopy ^110^ and drug delivery.^111^ Although protein denaturation is well known to occur on naked gold surfaces, predicting the resulting denatured protein structures remains a challenge.^112,113^ In this section, we demonstrate how to use DiffPIE to predict possible structures of proteins that bind to a gold surface. The system we study is amyloid *β*_16_*_−_*_22_ (A*β*_16_*_−_*_22_) with the KLVFFAE sequence on the surface of Au(111) (Fig. 3(a)), which has been used as a model system to study the environmental impacts on protein fibrillation.^114^ Additionally, the small size of this system allows us to exhaustively sample all possible conformations with metadynamics to validate DiffPIE. The biasing force *F*_0_(**x**) in this example is the thermodynamic force associated with binding A*β*_16_*_−_*_22_ to the gold surface. The force is approximated by the contributions of each residue. We perform metadynamics simulations of a dipeptide molecule for each type of amino acid in A*β*_16_*_−_*_22_ to generate a free energy surface (FES) based on three distances, *d*_1_, *d*_2_, and *d*_3_, as shown in Fig. 3(b). The detailed description of the construction of *F*_0_(**x**) will be presented in the “Methods” section. For residue *i* in A*β*_16_*_−_*_22_, we generate the force on its C*α*, while the C*α*–surface distances of residue (*i* − 1) and residue (*i* + 1) match *d*_1_ and *d*_3_, respectively. The design of *F*_0_(**x**) highlights the importance of the orientation of the residue, as reflected by the different values of *d*_1_ and *d*_3_. Unlike the previous example, **F**_0_ in this case cannot be expressed as the gradient of an energy function, highlighting the flexibility of designing **F**_0_. Similarly to the previous example, we project the structures generated by DiffPIE onto UMAP features trained from the benchmark metadynamics simulation. As shown in Fig. 3(c), the clusters of structures generated by DiffPIE overlap well with low-free-energy regions on the FES of metadynamics, suggesting that the populations of structures generated by DiffPIE are qual-itatively consistent with conformation stability. DiffPIE is also capable of finding structures similar to representative conformations from metadynamics simulations. We emphasize that while some configurations are highly flexible and DiffPIE can only recover structures with similar overall shapes, for conformations that are energetically stabilized with low free en-ergy, DiffPIE can generate structures that nearly overlap with the simulated structures. Due to strong interactions between amino acid and gold surfaces, these representative structures are not sampled by the original Str2Str algorithm (see Supporting Information). Again, generating structures with DiffPIE only took less than one hour, while metadynamics simu-lations may take days to complete. Finally, we tested the relaxation of generated structures by randomly selecting three structures as starting points for short MD simulations. Both the starting structures and the relaxation trajectories are presented in Fig. 3(d). For some starting structures, the relaxation leads to the same final conformation, suggesting that rep-resentative structures of stable conformations are included in our predictions. Meanwhile, high-free-energy structures quickly relax, indicating that even short MD simulations can significantly aid structure refinement.

**Figure 3:**
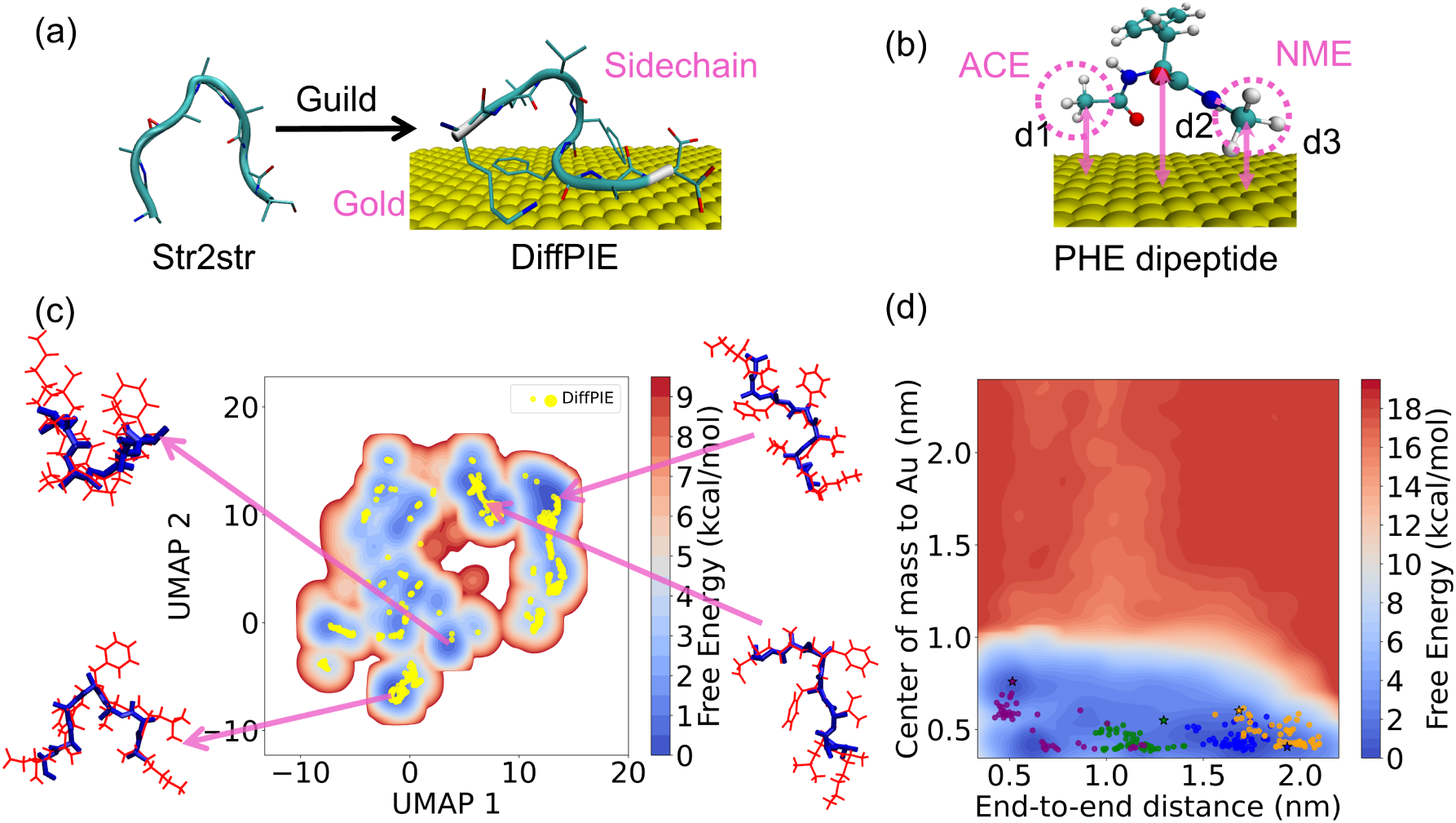
(a) presents the system of *Aβ*_16_*_−_*_22_ on Au(111) surface. (b) demonstrates a model dipeptide molecule (PHE in this case) binding on the gold surface to construct the bias-ing force **F**_0_. The binding free energy along three CVs, *d*_1_, *d*_2_, and *d*_3_, was evaluated by a metadynamics simulation. Here, *d*_1_, *d*_2_, and *d*_3_ are defined as the distances from the gold surface to the methyl carbon of the acetyl cap, the *α*-carbon, and the methyl carbon of the N-methylamide cap, respectively. (c) shows the comparison between metadynamics simulations and DiffPIE. The background FES was generated by projecting the metady-namics simulation trajectory onto UMAP features. The DiffPIE samples are projected onto the same UMAP feature space as yellow dots. We further present the top-down view of representative simulated structures (red) from low free energy clusters and similar DiffPIE samples (blue), with the gold surface oriented parallel to the plane of the paper. (d) shows short MD relaxations of three structures selected randomly from DiffPIE structure clusters. The starting structures (stars) and their relaxation trajectories (dots) are mapped onto the FES of the *Aβ*_16_*_−_*_22_ end-to-end distance and the *Aβ*_16_*_−_*_22_ center-to-gold surface distance.

## Discussion and Conclusion

The two examples presented in this study demonstrate that the introduction of PEI into the foundation generative model (DiffPIE) is capable of exploring protein-with-PEI conformations, which is an impossible mission with the foundation generative model. We highlight that DiffPIE does not require any further training or fine-tuning of the foundation generative model. In both examples, our objective was to achieve a trade-off between the accuracy of PEI models and the computational cost of building them. Thus, approximations were used in both cases. In the first example, we ignored the nonbonded interactions between the organic linker and residues other than cysteines. In the second example, the presence of the gold surface disrupts the solvation shell of residues near the surface, leading to changes in the solvation free energy that are non-additive. The generated structures suggest that such approximations do not significantly affect the downgrade quality of structures generated by DiffPIE, which suggests that DiffPIE does not require an exact PEI model for the generation of peptide structures. Moreover, different designs of the biasing force in two examples highlight the flexibility of designing PEI models in DiffPIE. The generated structures can also be used as starting points for MD simulations or enhanced sampling simulations.^115,116^ As suggested in recent studies, the introduction of enhanced sampling simulations starting from multiple important conformations predicted by machine learning models can significantly improve sampling efficiency.^117,118^ We expect that DiffPIE can be combined with enhanced sampling methods or other free-energy calculation approaches to further improve structural exploration and thermodynamic analysis.

Ideally, we hope that DiffPIE will generate structures following the Boltzmann distribution of a peptide with PEI. However, in practice, it is limited by several theoretical and technical challenges. First, generating structures that follow the Boltzmann distribution of a protein requires that the foundation generative model be able to generate protein structures following the Boltzmann distribution without PEI. The foundation generative model used in this study, Str2Str, is known to be biased towards generating more helix-like structures. As shown in the “Results” section, the bias force in DiffPIE can help correct this bias but cannot fully eliminate it. Some recent studies have proposed methods to correct Str2Str^95^ or to develop new foundation generative models capable of generating peptide ensembles without PEI.^119,120^ We expect that improved foundation generative models will also enhance the performance of DiffPIE. Second, DiffPIE is limited by the accuracy of the PEI models. Therefore, we do not expect the current version of DiffPIE to generate structures that fully follow the Boltzmann distribution of proteins with PEI. Improving the quality of the DiffPIE-generated ensemble remains an important goal for future work.

Another challenge in DiffPIE is how to efficiently identify structures that work best as initial structures for sampling. The current solution is to screen structures using short MD simulations, which provides the most reliable, but also the most computationally expensive, screening approach. Other methods, such as energy minimization in vacuum or with an implicit solvent, work well for the cyclic peptide example but perform poorly for the peptide-on-gold example. Alternative energy scoring functions for proteins, such as the Rosetta energy functions,^121^ are typically tuned for proteins without PEI and are not directly applicable to either example without careful reparameterization. Therefore, the development of efficient and accurate screening methods for proteins with PEI remains an important area for future research.

In summary, we have developed DiffPIE, a framework that guides a foundation diffusion model with physics-based interactions to generate protein structures with PEI. This framework does not require additional training and can be adapted to various types of PEI. In addition to the examples discussed in this work, DiffPIE has the potential to greatly broaden the applications of protein foundation diffusion models, including protein–ligand binding, proteins at interfaces, and proteins in external fields.

## Methods

Our proposed method is general for any score-based diffusion model (SBDM). A SBDM perturbs data distribution to noise prior with a diffusion process over a unit time by a linear stochastic differential equation (SDE):

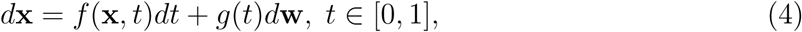

where *f* (**x**, *t*), *g*(*t*) are user-defined drift and diffusion functions of the SDE and **w** denotes a standard Wiener process. With a carefully designed SDE, the marginal probability of **x** at diffusion time *t*, *P_t_*(**x**), changes from the data distribution *P* (**x**) to approximately a simple Gaussian distribution. In this paper, we use an SDE with drift term *f* (**x**, *t*) = 0 and diffusion function 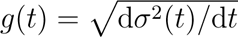, where *σ*(*t*) represents the noise scale. For any diffusion process in Eq.(4), it has a corresponding reverse-time SDE^99,122^ :

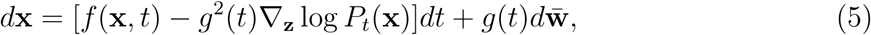

with **w̅** a standard Wiener process in reverse time. The trajectories of the reverse SDE have the same marginal densities as those of the forward SDE. Thus, the reverse-time SDE can gradually convert noise to data. The SBDM parameterizes the time-dependent score function ∇**_x_** log *P_t_*(**x**) in reverse SDE with a neural network **s*_θ_***(**x**(*t*), *t*).

Suppose that the SBDM baseline aims to predict protein structures that follow the Boltzmann distribution, i.e. *P* (**x**) ≈ *p*^NPIE^B(**x**) ∝ exp{−*βU* (**x**)} where *U* (**x**) is a protein energy function that is similar to a “statistical potential”. *β* = 1*/k*_B_*T* where *k*_B_ is the Boltzmann constant and *T* is temperature. PEI can be written as an extra energy term *U*_ext_(**x**). The Boltzmann distribution of protein structures with PEI is 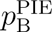(**x**) ∝ exp{−*β*(*U* (**x**)+*U*_ext_(**x**))} It has been proven that sampling *P*_bias_(**x**) ≈ 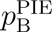(**x**) requires a reverse diffusion process with time-dependent marginal distribution satisfying: *P*_bias_*_,t_*(**x**(*t*)) ∝ *P_t_*(**x**(*t*)) exp{−*βU_t_*(**x**(*t*))} where *U_t_* is a time-dependent energy function defined as^95^

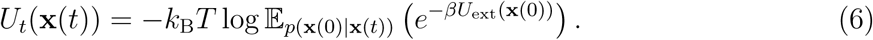

*p*(**x**(0)|**x**(*t*)) in Eq.(6) is the probability of **x**(0) conditional on the location of **x**(*t*) in the unmodified revert diffusion process. To further simplify the expression of *U_t_*, we adopted an approximation used in the manifold constraint algorithm for the diffusion model:

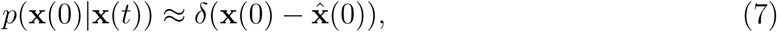

where *δ*(·) is the Dirac delta function and **x̂**(0) = E(**x**(0)|**x**(*t*)). In certain types of diffusion models such as DDPM, **x̂**_0_ is an explicit function of **x**(*t*) and *t*: **x̂**_0_ = **h**(**x**(*t*), *t*).^98^ With the approximated *p*(**x**(0)|**x**(*t*), Eq.(6) can be simplified to *U_t_*(**x**(*t*)) ≈ *U*_ext_(**h**(**x**(*t*), *t*)). The approximation of using Eq.(7) works better when *t* is close to 0. Therefore, care must be taken at the initial stage of backward diffusion. In this study, we scaled **F**(**x**(*t*), *t*) by a time-dependent factor *λ*(*t*). *λ*(*t*) is closed to 0 when *t* is closed to 1 while *λ*(*t*) increases to 1 when *t* approaches 0. The detailed designs of *λ*(*t*) are presented in the supporting information. With *U*_ext_(**x̂**(0)), we can calculate the potential *U*_ext_ and we can push the potential pack to the diffusion model: **F**(**x**(*t*), *t*) = −∇**_x_**_(_*_t_*_)_*U*_ext_(**h**(**x**(*t*), *t*)). Although using *U*_ext_ provides a clear picture of how to connect DiffPIE to the Boltzmann distribution of the protein with PEI. We can relax the requirement of using an interaction potential *U*_ext_ for the structure generation task. This provides a more flexible tool for users to design **F**, such as **F** representing non-equilibrium perturbations from the environment. Therefore, in general, we will express **F** = **F**[*U*_ext_](**x**(*t*), *t*) to highlight the dependency of **F** on *U*_ext_.

In this work, two examples of **F** are used: one with an explicitly defined **F**(**x**(*t*), *t*) = −∇**_x_***_(t)_U*_ext_(**h**(**x**(*t*), *t*)), and another that uses **F** depending on *U*_ext_ in a more complicate way, as describe later in this paragraph. In the example of the cyclic peptide P3-F, we use the FES of six distances (Fig. 2(b)) as *U*_ext_. This six-dimensional potential was fitted using a logarithmic function of a mixture of Gaussians. We emphasize that the selected collective variables (CVs) provide joint information on both the distance and orientation between the two cysteines. In the second example, **F** is designed as follows. First, we use metadynamics to generate a binding FES, *A_i_*, for a dipeptide molecule that represents the residue *i* of the protein on the gold surface. *A_i_*is a function of three distances, pre-sented in Fig. 3(b), and can be viewed as a function of **y**_1_, **y**_2_, and **y**_3_, representing the C*α* atom (**y**_2_) and the two methyl carbons (**y**_1_ and **y**_3_) of the two capping groups. Therefore, we can express *A_i_* as *A_i_*(**y**_1_, **y**_2_, **y**_3_). We define the biasing drift on the *i*-th C*_α_*, **F**^(^*^i^*^)^, as 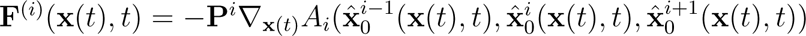 where 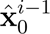, 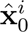, and 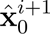 are the coordinates of the (*i* − 1)-th, *i*-th, and (*i* + 1)-th C*α* atoms in **x̂**_0_, respectively. **P***^i^* is a projection operator that only keeps ∇**_x_**_(_*_t_*_)_*A_i_* on the the *i*-th C*_α_*. Please refer to the Supporting Information for details on the generation of **F**.

Str2Str builds protein structures by applying translations and rotations in residue-specific local frames. Biasing forces are applied only to C*α* Cartesian coordinates, not orientations, since Tweedie’s formula does not extend to *SO*(3),^104^ and projecting forces onto the *SO*(3) tangent space can cause numerical instability.^95^ Generating side chain conformations consistent with both the backbone structure and PEI is essential, as side-chain-environment interactions strongly influence the binding affinity to material surfaces. Str2Str provides only backbone atoms, so side chain backmapping typically uses FASPR.^123^ However, FASPR alone is insufficient under PEI, as it can create steric clashes with surfaces or cannot build the required linkers. To overcome this, we developed an algorithm to reconstruct missing components (side chains and linkers) from a backbone structure *A* generated by DiffPIE, which corresponds to MD-derived structures used in *U*_ext_. In the first example, the P3-F CYS side chains and the organic linker were reconstructed by aligning the backbone atoms in *A* with MD snapshots. In the second, the side chains near the gold surface were rebuilt by selecting dipeptide conformations from MD that (1) align with the backbone frame of *A* and (2) match residue–surface displacement. The remaining side chains were completed using FASPR.

## Acknowledgement

Y.W. and M.C. gratefully acknowledge the support of the American Chemical Society Petroleum Research Fund (Grant Number 67307).

## Supporting Information Available

All codes and scripts used in this study are available at [GitHub (DiffPIE)](https://github.com/mingchenPurdueChem/DiffPIE). The molecular dynamics (MD) trajectories and generated results can be accessed at [Google Drive](https://drive.google.com/drive/folders/1dFnwRA-LjiuNRPm_x5LmsAWaiW4Khl0h?usp=sharing).

The following files are available free of charge.

- Supporting information — Extrapolating Foundation Generative Models with Physics: a Case Study of Exploring Peptide Conformations under Protein-environment Inter-actions: addtional details of biasing force preparation, simulation setup, and DiffPIE structure generations.

## TOC Graphic

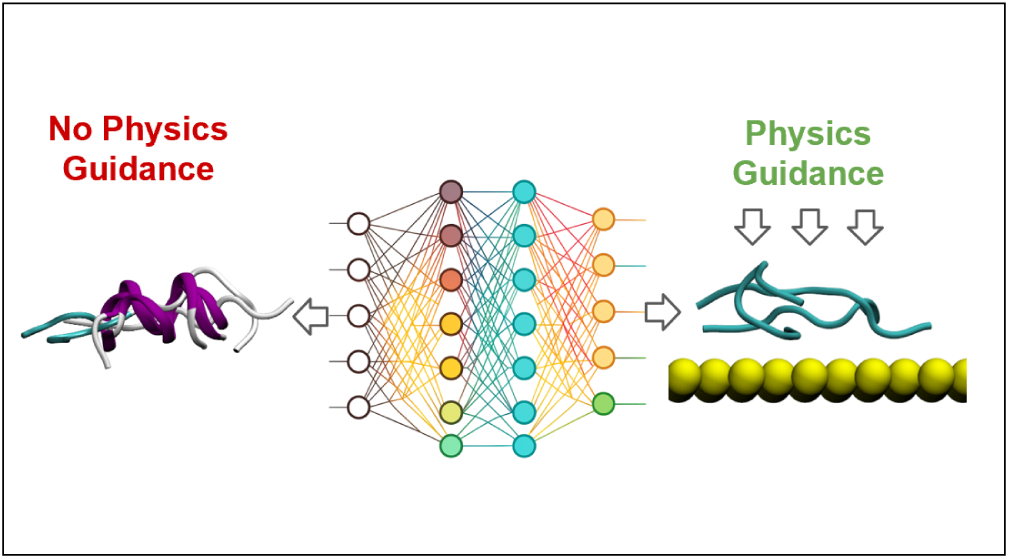

