## Supplementary Information for "DiffPIE: Guiding Deep Generative Models to Explore Protein Conformations under External Interactions"

### Details of Generating Biasing Force

Biasing forces were constructed using specifically designed model systems. In the first example, the sequence of the peptide is CDPETGECL. The stapler molecule is linked to the peptide through C-S bonds of the two cysteines. The biasing force acts on the peptide without the stapler molecule to “mimic” the impact of the presence of the stapler molecule. To obtain the biasing force, we built a model system described as follows: first, we built two cysteine dipeptides with acetyl and N-methylamide capping groups; next, the stapler molecule was linked to the side chains of two cysteine dipeptide molecules. Molecular dynamics (MD) simulations were performed to sample the configurations of the model system in aqueous solution. The force field parameters for the stapler molecule were generated using GAFF2,[1] and the system was solvated with 1228 SPC water molecules. Equilibration was conducted by energy minimization followed by a 20 ps NVT simulation at 300 K using the V-rescale thermostat with a coupling constant of 0.1 ps. All bonds involving hydrogen atoms were constrained using the LINCS algorithm. The integration was performed using the velocity-Verlet algorithm with a 2 fs timestep. Long-range electrostatics were treated with PME-Switch, with a real-space cutoff of 1.2 nm and a switching function starting at 1.16 nm. Van der Waals interactions were computed with a cutoff scheme and a potential-shift-Verlet modifier over the same range. Production sampling was carried out using a 100 ns NPT simulation at 300 K and 1 bar using isotropic Parrinello–Rahman pressure coupling with a time constant of 2.0 ps. The simulations were prepared and performed using GROMACS, version 2022.5.[2] Six distances were selected as collective variables (CVs) to describe the configurational change of the model system, as defined in Fig.2(b). We built the probability density $P$ by fitting the CV distribution with the Gaussian mixture model(gmm) with 10 Gaussians. $P$ was then converted to a biasing energy $-k_{\mathrm{B}}T \log P$ where $k_{\mathrm{B}}$ is the Boltzmann distribution and the temperature $T$ is 300 K. Here, we display the 2D projection of the six distances in Fig. S1, Fig. S2, and Fig. S3.

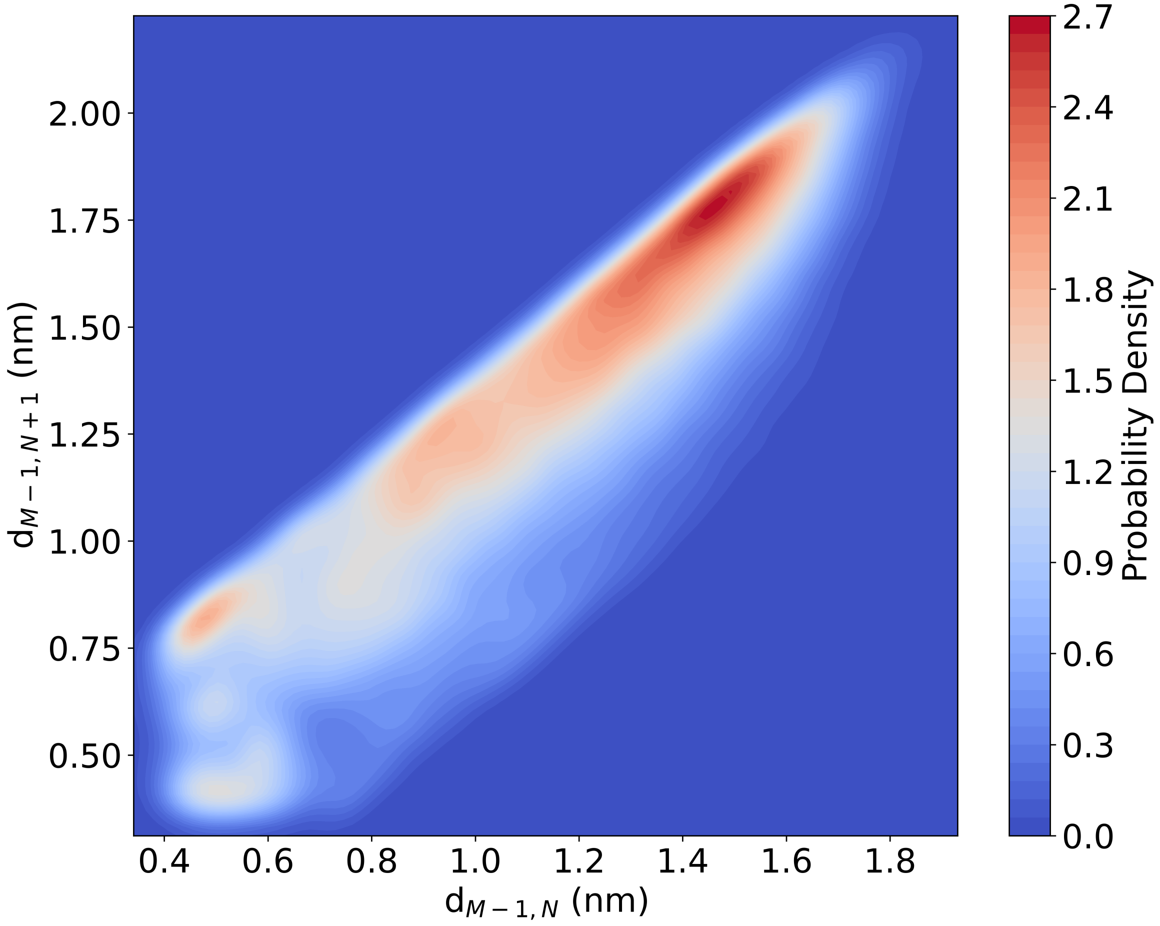


Figure S1: The probability density of $d_{M-1,N}$ and $d_{M-1,N+1}$.

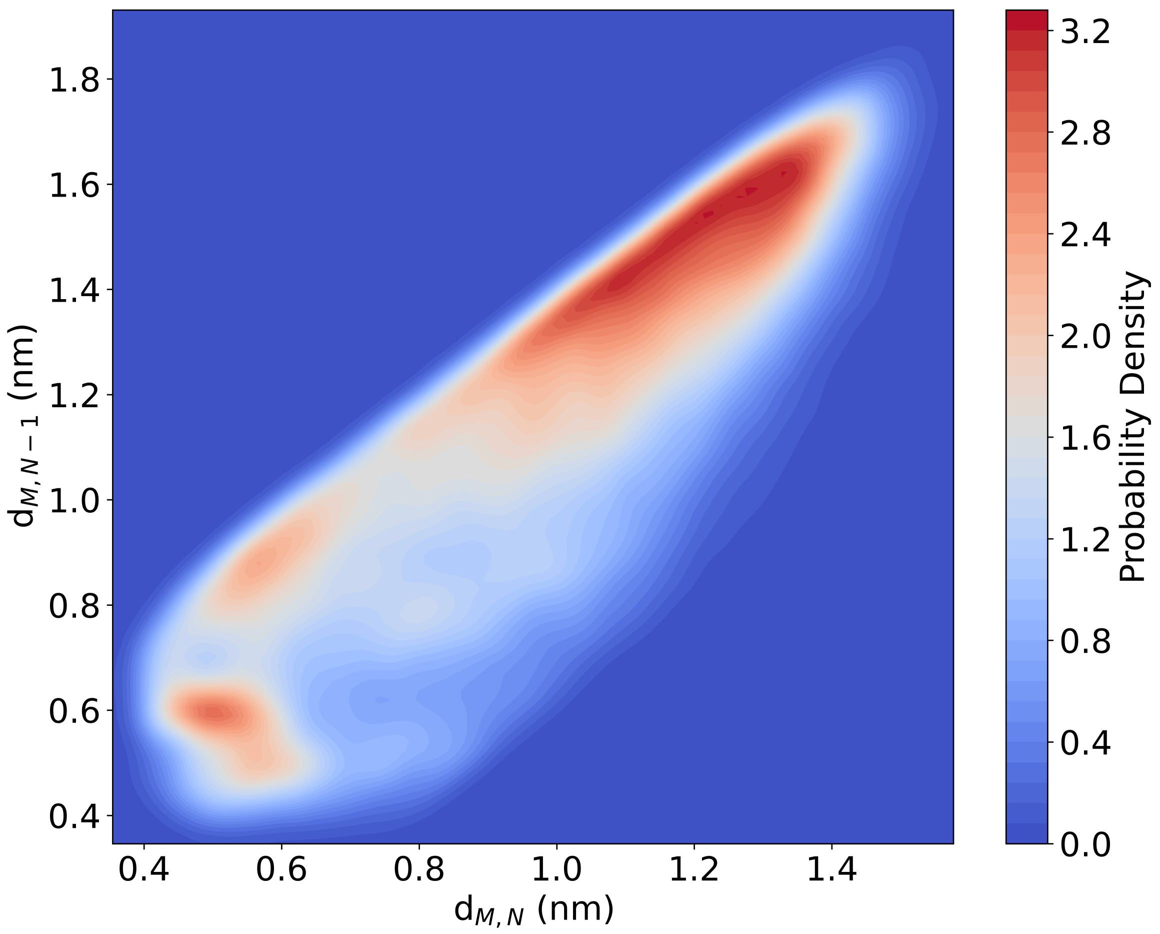


Figure S2: The probability density of $d_{M,N}$ and $d_{M,N-1}$.

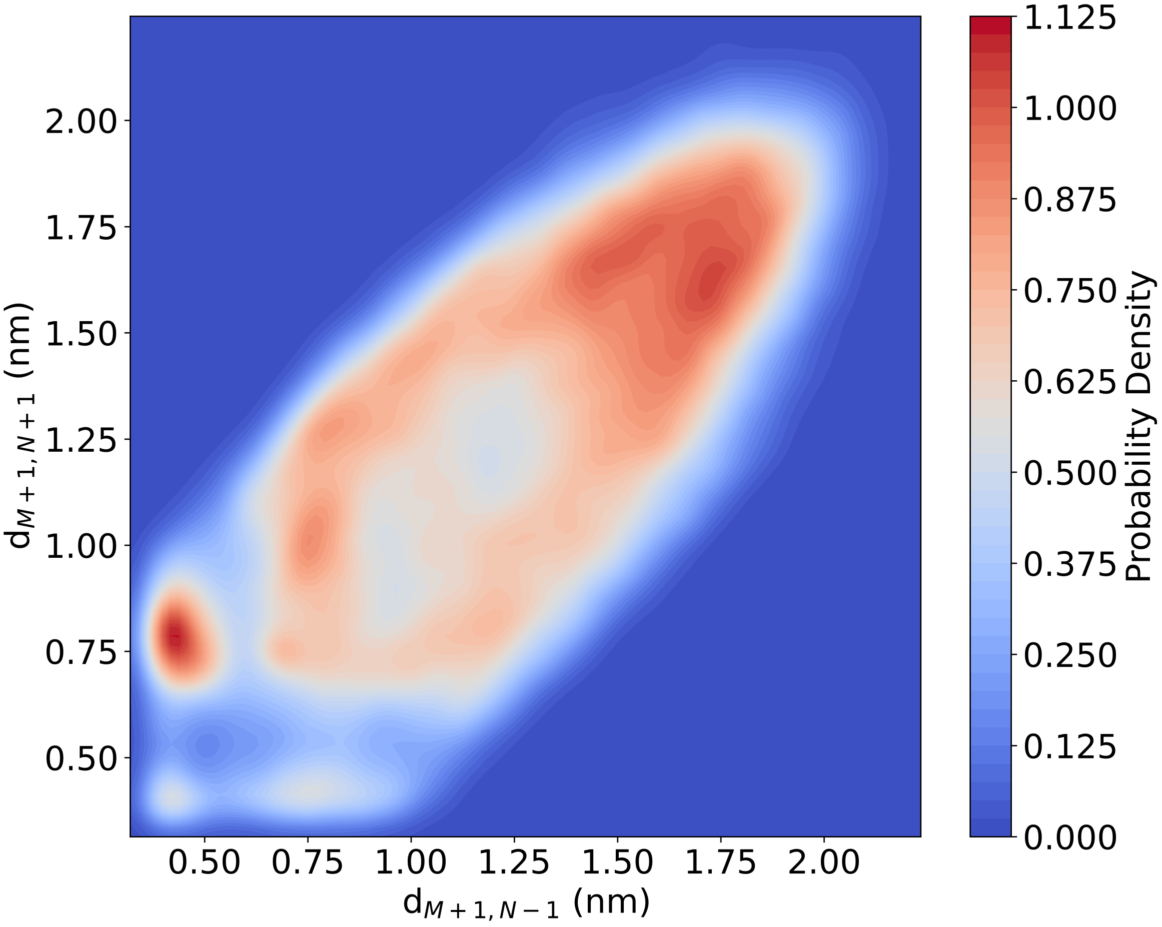


Figure S3: The probability density of $d_{M+1,N-1}$ and $d_{M+1,N+1}$.

In the second case, the sequence of the peptide is KLVFFAE. We prepared the dipeptide model for each type of amino acid to describe the restraints defined in the main content. We used the GoldP force field to simulate the Au(111) surface and OPLS-AA for the peptide parameters.[3,4] We set five layers of Au with ABC stacking. Since the adsorption of the peptide is strong, we employed well-tempered metadynamics simulations to accelerate the sampling of configurations (shown in Fig.S4). The system was initially put into an orthorhombic box of 4.1 $\times$ 4.1 $\times$ 5.0 $nm^3$. It was solvated using SPC water, resulting in a total of 1910 to 1916 water molecules, depending on the amino acid type. The system was equilibrated by energy minimization, followed by a 200 ps NVT simulation and a 200 ps NPT simulation. Production sampling was conducted in the NVT ensemble. Well-tempered metadynamics simulations were performed using GROMACS 2022.5 patched with PLUMED 2.9.[5] A CV was defined as the distance between the C$\alpha$ atom and the gold surface. Gaussian functions were deposited every 500 steps, with an initial Gaussian height of 1.5 kJ/mol and a Gaussian width ($\sigma$) of 0.01 nm. The bias factor was set to 6.0, and the reference temperature was 300 K. All simulations were unbiased to recover the Boltzmann distributions.[6] Then we used the three distances defined in Fig.3(b) in the main article as CVs to build the free energy surface, from which the biasing force was further constructed, as discussed in the main article. Similarly to the first example, the projection of $-k_{\mathrm{B}}T \log P$ of C$\alpha$ onto the gold distance, one of the three defined distances, is plotted for each residual(Fig.S5).

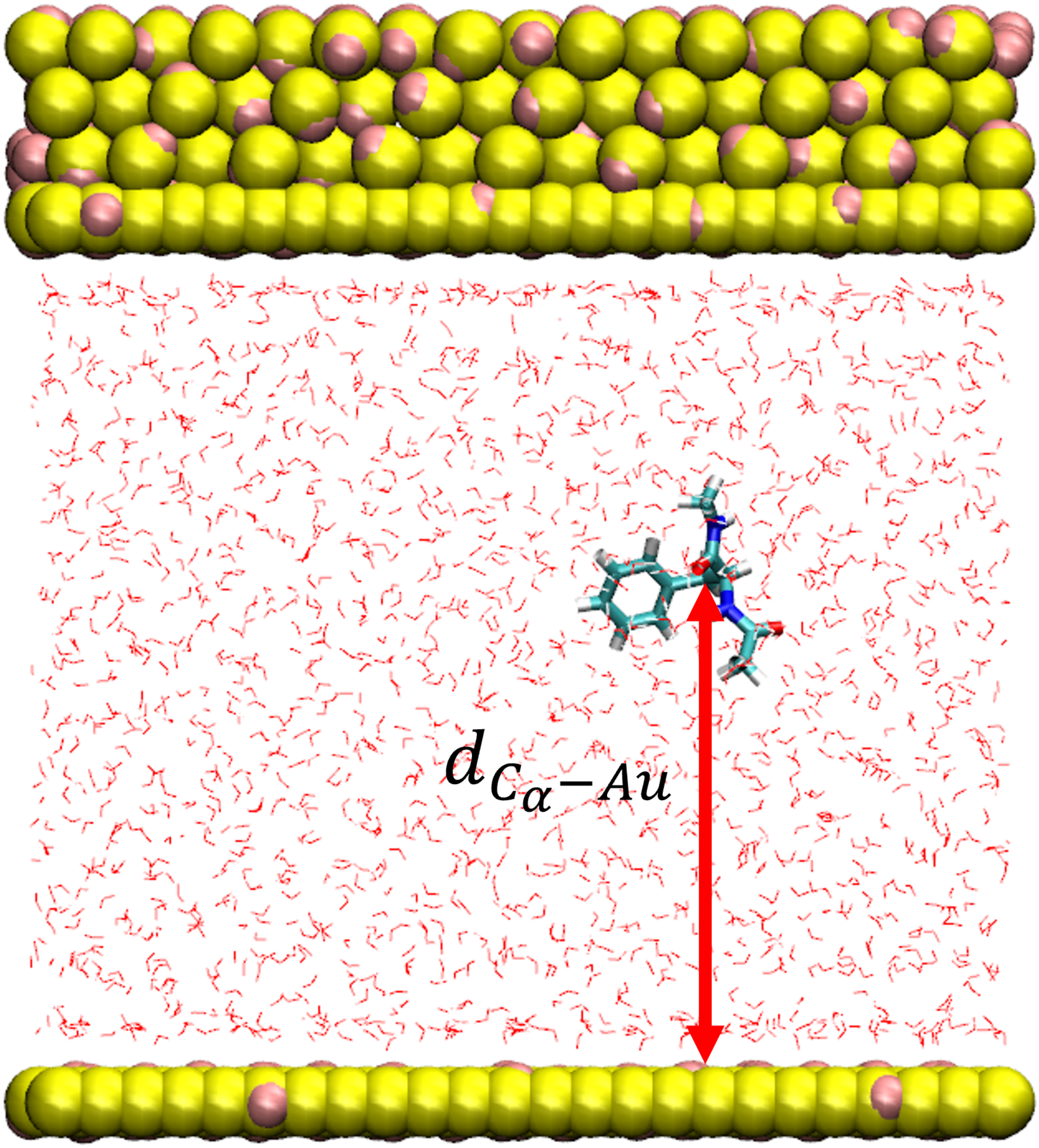


Figure S4: The metadynamics simulation setup for biasing force generation with a PHE dipeptide model. Distance between C$\alpha$ and Au surface is used as CV. The system is solvated by SPC water. The yellow particles are fixed gold, and the pink particles are the image charge particles. Visualized by VMD.[7]

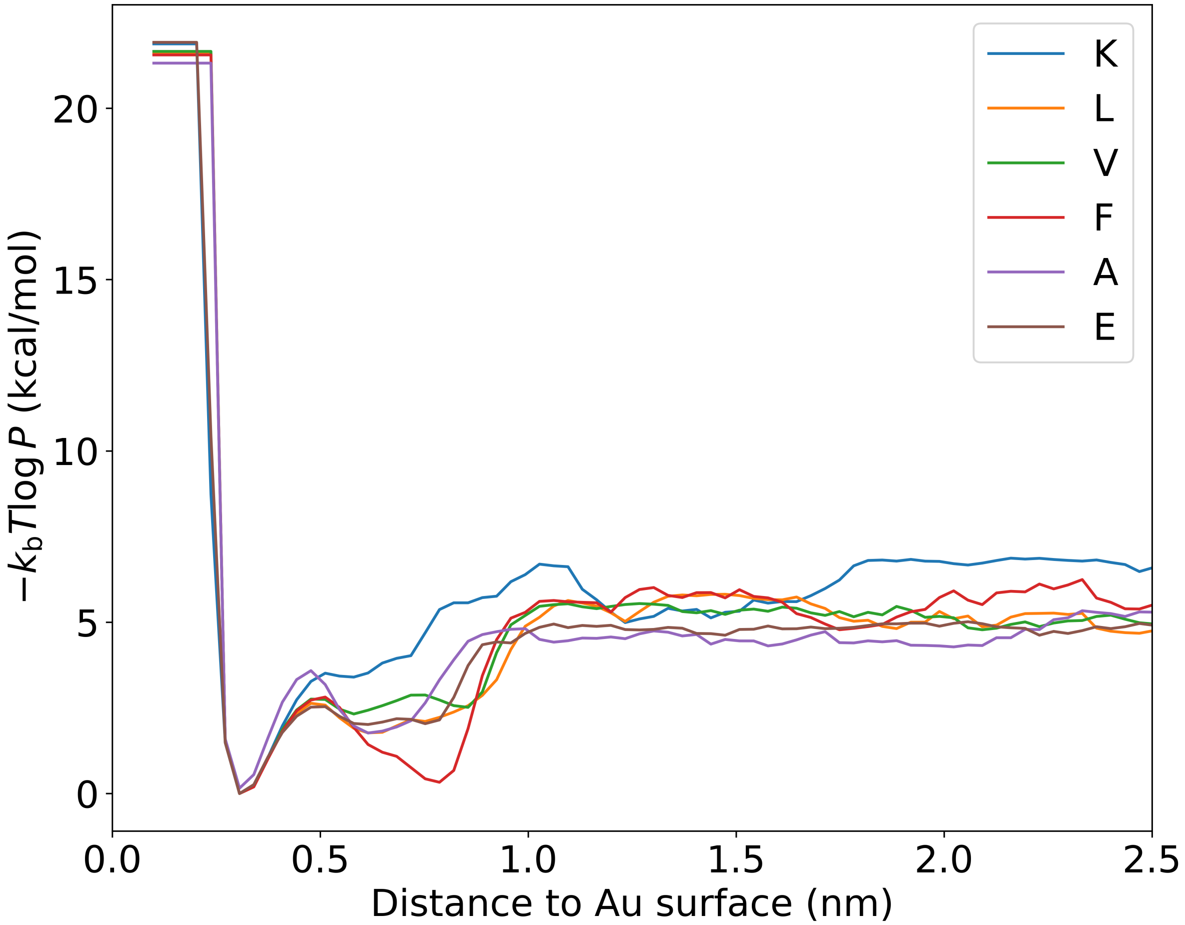


Figure S5: The biasing energy of all the dipeptide models used to construct biasing force as a function of $C_\alpha$ to Au distance.

When the biasing force is applied to the diffusion model, we adopt an approximation that $p(\mathbf{x}(0)|\mathbf{x}(t)) \approx \delta(\mathbf{x}(0) - \hat{\mathbf{x}}(0))$ where $\mathbf{x}(t)$ is the diffusion variable at time $t$. $\hat{\mathbf{x}}(0)$ is the expectation of $\mathbf{x}(0)$ conditional on $\mathbf{x}(t)$. This approximation is less accurate when $t$ is close to 1. Here, we used a function, force_scale, that gradually changes $\eta$ from 0 to 1 to ensure that the biasing drift near the end of the reverse diffusion process ($t = 0$) is $\eta \mathbf{F}(\mathbf{x}, t)$ while the biasing drift at the beginning time of the reverse diffusion process is approximately 0. Many functional forms can be used for this purpose. Here, we employ (S.1) for the case of a cyclic peptide and (S.2) for the case of a peptide on gold:

$$\text{force\_scale}(t) = \frac{1}{1 + e^{-\beta(t_{\text{mid}} - t)}}, \quad t_{\text{mid}} = 0.3 \tag{S.1}$$

$$\text{force\_scale}(t) = 1 - \frac{e^{mt} - 1}{e^{m} - 1}, \quad m = 1.5, \quad 0 \leq t \leq 1 \tag{S.2}$$

### Details of Benchmark Metadynamics Simulations

To explore the free energy landscape of the full system, we performed well-tempered metadynamics to generate conformations as the ground truth. In the first case, we sampled the Apo state of P3-F, using the end-to-end distance as the CV, and the system was solvated with 3322 SPC water molecules. In the second case, we investigated the adsorption of $A\beta_{16\text{–}22}$ onto a gold surface. The box size was the same as the one used in the previous section and the system was solvated with 1870 SPC water molecules. We used two CVs: the COM-to–Au distance and the end-to-end distance. Both systems underwent energy minimization, followed by 200 ps of NVT equilibration and 200 ps of NPT equilibration. Production sampling was then carried out in the NVT ensemble. Metadynamics parameters were identical to those described in the previous section.

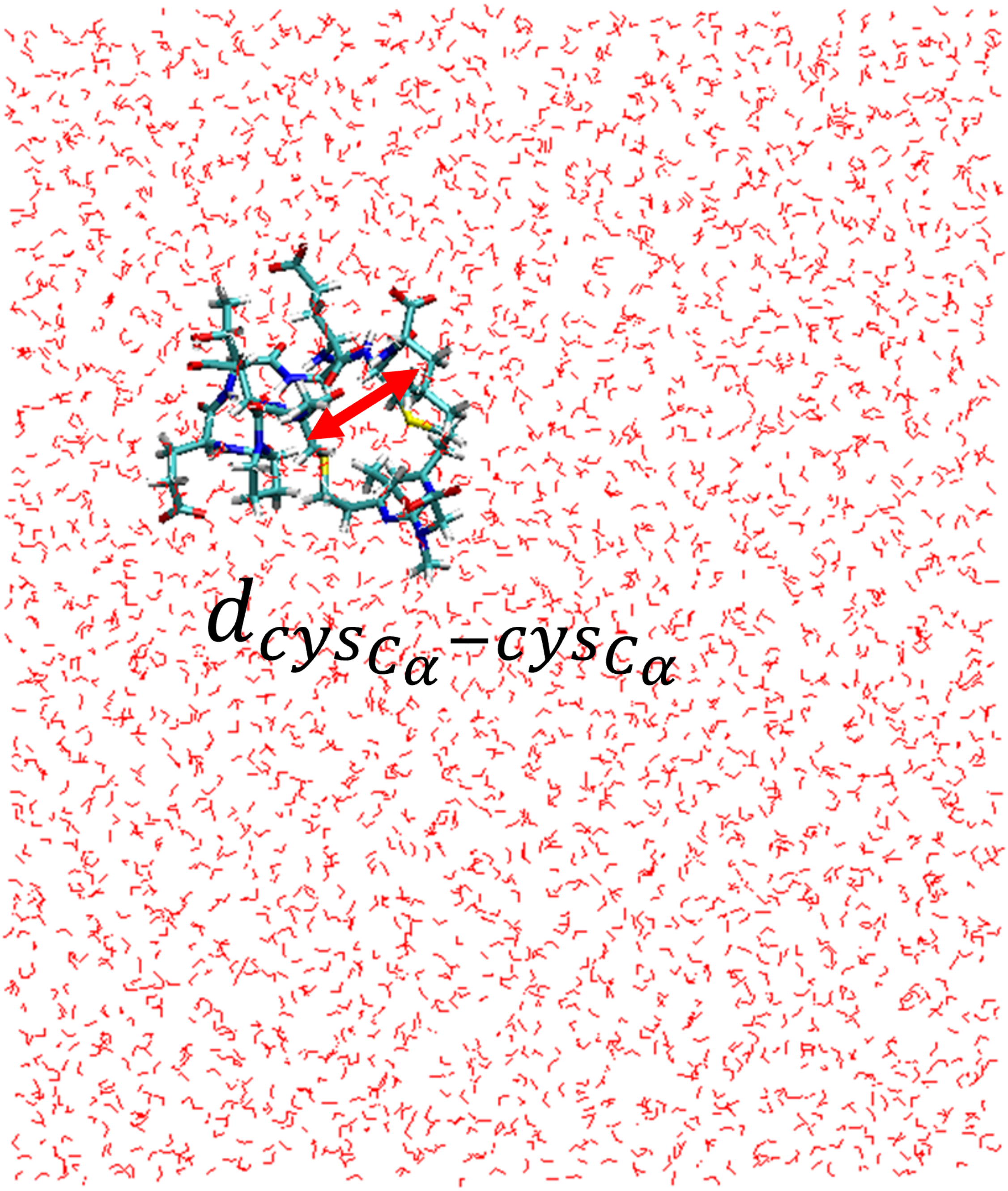


Figure S6: The setup of metadynamics sampling of P3-F Apo state using distance between two $C_\alpha$ of cysteines as CV.

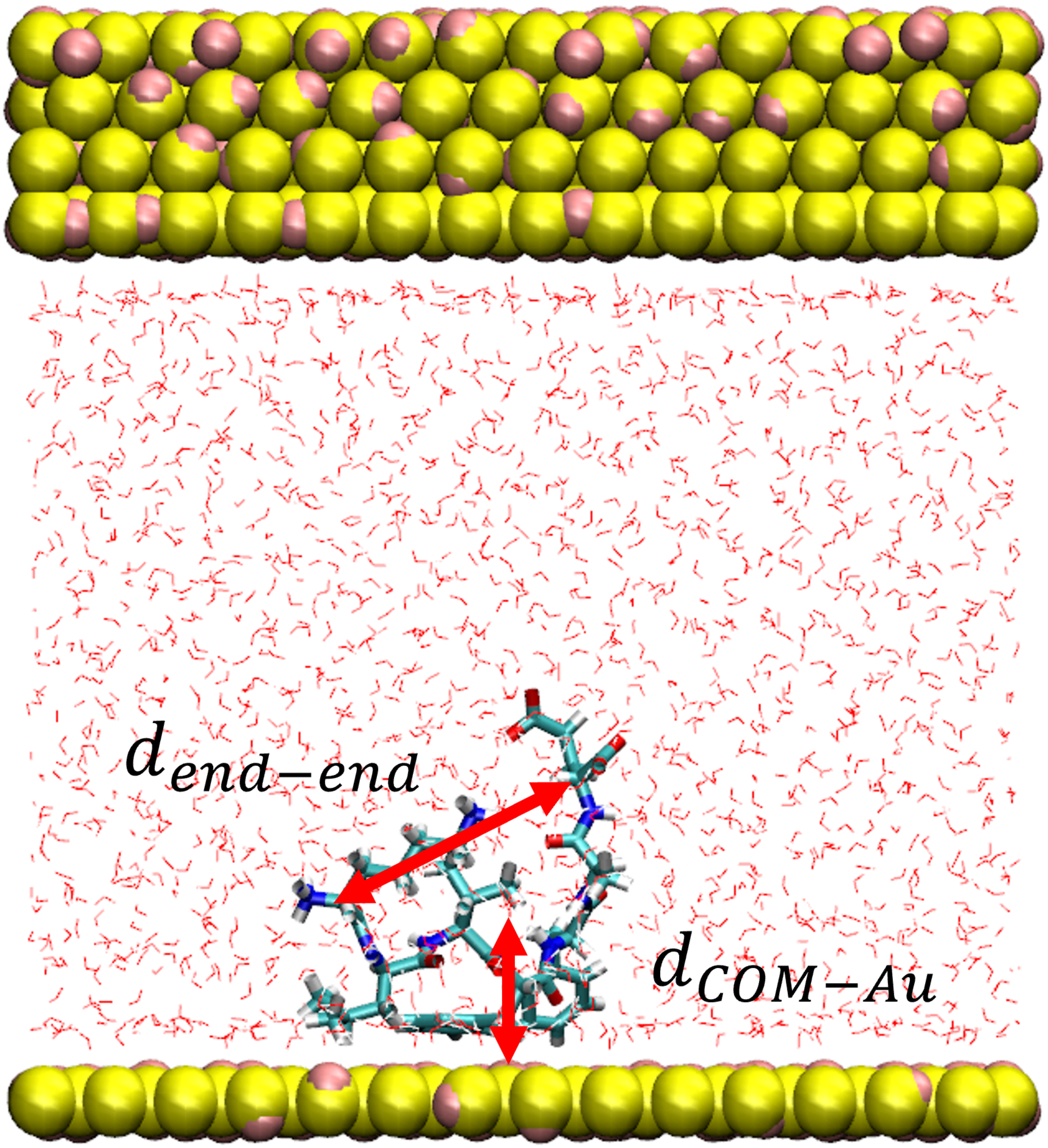


Figure S7: The setup of metadynamics sampling of $A\beta_{16-22}$ on Au111 surface using end-to-end and center of mass to gold distances as CVs.

#### Details of DiffPIE Structure Generations

One challenge with the use of Str2Str is how to properly apply the biasing force to terminal residues. We used C$\alpha$ of residue $i-1$ and C$\alpha$ of residue $i+1$ to determine the orientation guidance for residue $i$. There is no C$\alpha$ of residue $i-1$ if residue $i$ is N-terminus and there is no C$\alpha$ of residue $i+1$ if residue $i$ is C-terminus. To solve the problem, we capped the terminus with one more residue, typically glycine or alanine. For P3-F pepetide, the N-terminus cysteine was capped with an alanine residue to allow the application of six distance restrictions. For $A\beta_{16\text{–}22}$, both the N-terminus and C-terminus were capped with glycine to serve the same purpose. The capped residues will be removed in the post-analysis steps. We compared the structural ensemble generated with capped and uncapped sequences using Str2Str to determine whether a glycine or alanine cap should be used, based on the degree to which the generated structural ensemble is perturbed by cap residues. We emphasize that similar approaches, such as linking two proteins with a glycine chain, have been used in some protein structural generation models such as ESMFold.[8,9]

The strength scaling constant of the biasing force was chosen empirically. In principle, numerical stability during the reverse diffusion process, which favors small perturbations, must balance with the enforcing structural bias on the generated conformations, which benefits from a stronger bias. Users may adjust this rescaling constant based on specific use cases.

During inference using DiffPIE for P3-F pepetide, we initialized the process using the structure predicted by AlphaFold3 and set Str2Str to operate in reverse-only mode. A second generation iteration was then initiated using a structure with a loop conformation from the first iteration, since Str2Str tends to generate undesired helical structures (as discussed in the main article). In this case, we started the reverse diffusion process at time $t = 0.3$. Starting from other loop structures generates similar results. For $A\beta_{16\text{-}22}$ adsorption on the gold surface, we also used the predicted structure of AlphaFold3 as a starting point. A harmonic repulsive potential centered at 0.1 nm away from the Au surface was applied to prevent unwanted atom clashes. Initially, the Au surface was placed 0.9 nm away from the peptide's center of mass (COM). This distance was gradually reduced to a final value of 0.2 nm during the reverse diffusion process, slightly below the low free-energy basins identified in metadynamics simulations.

#### Details of Side Chain Generations and Structure Assembly for P3-F

We first generated side chain coordinates for DiffPIE generated structures using FASPR and GROMACS.[10] To allow structural alignment, both the stapler molecule and the peptide were transformed into a new local coordinate frame defined by the N, C$\alpha$, and C atoms of the first cysteine residue, with the C$\alpha$ atom set as the origin. After this transformation, the two structures were assembled according to two criteria: (1) absence of steric clashes between the peptide and the organic stapler molecule (atomic distance greater than 1 Å) and (2) formation of a C-S bond of physically reasonable length (no longer than 2.2 Å).

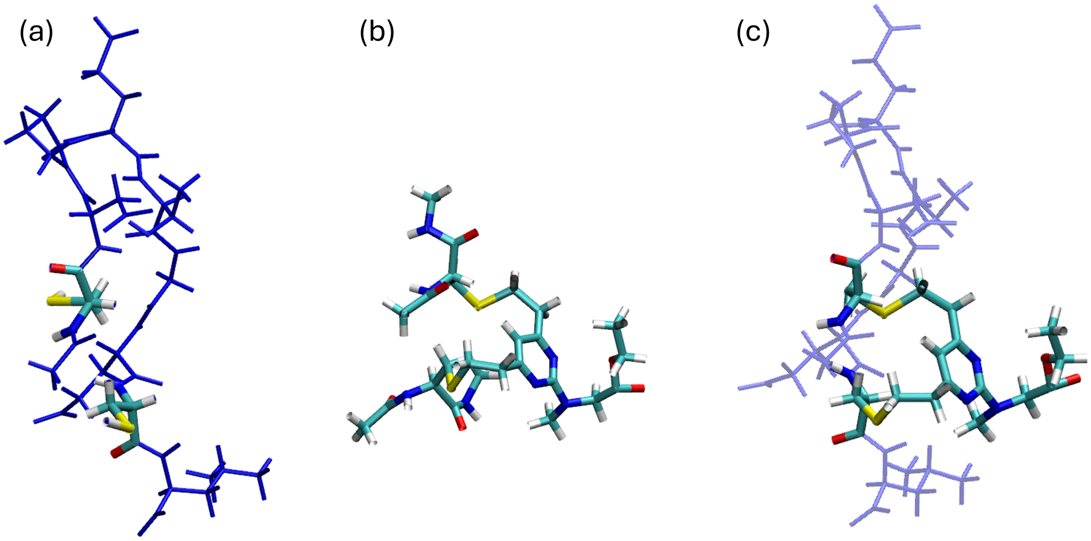


Figure S8: Assembly of P3-F from DiffPIE. (a) A structure with backbone sampled by DiffPIE, side chain generated by FASPR, and H generated by GROMACS. (b) Stapler molecule sampled by MD simulations, selected by matching the C$\alpha$ coordinates of cysteines with these in the structure in (a). (c) The assembled structure with cysteines side chain replaced structure in (b).

### Details of COM-to-Au adjustment for $A\beta_{16-22}$

The structures generated near the Au surface by DiffPIE do not always adopt optimal adsorption positions. This is primarily due to the soft repulsive interactions between the peptide and the Au surface. To address this, we readjusted the COM position of the peptide relative to the Au surface by minimizing the total adsorption energy $E_{\mathrm{ab}}$. The total adsorption energy $E_{\mathrm{ab}}$ is approximate by the summation of the contributions of each residue $E_{\mathrm{ab}}^{(i)}$: $E_{\mathrm{ab}}(z_1, \cdots, z_N) = \sum_{i=1}^{N} E_{\mathrm{ab}}^{(i)}(z_i)$ where $z_i$ is the distance between C$\alpha$ of residue $i$ and the gold surface. $E_{\mathrm{ab}}^{(i)}$ was 1D binding FES that was generated from the metadynamics simulations performed for the preparation of $\mathbf{F}_0$ (as shown in Fig.S5). Importantly, only the global position of the peptide was adjusted as a rigid body, rather than the positions of individual residues. The position optimization results are illustrated in Fig. S9.

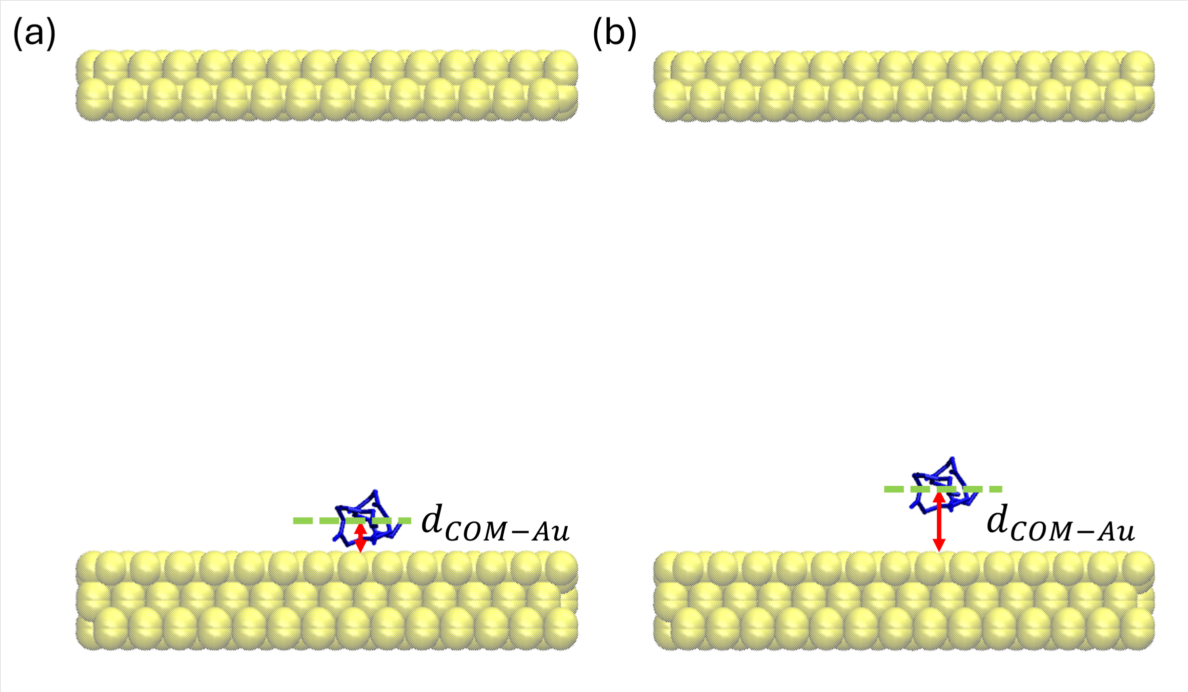


Figure S9: COM-to-Au adjustment. (a) A DiffPIE sample placed near Au surface with COM-to-Au distance initiated at 0.5 nm. (b) Adjusted COM-to-Au distance after restraint energy minimization.

#### Effectiveness of DiffPIE for $A\beta_{16-22}$

To demonstrate the effectiveness of DiffPIE in generating initial protein structures with proper interactions on Au(111), we also landed $A\beta_{16-22}$ structures generated from the bare Str2Str model on the gold surface using the method discussed in Section "Details of COM-to-Au adjustment for $A\beta_{16-22}$", and analyzed the distances of the first and last residues from the gold surface. The original Str2Str distribution exhibited two distinct clusters: when the N-terminus (first residue) was close to the surface, the C-terminus (last residue) tended to orient away from the surface, and vice versa. (Fig.S10(a)). In contrast, DiffPIE effectively modified this distribution, producing a single cluster where both terminal residues remained close to the surface at a moderate distance of approximately 0.5 nm (Fig.S10(b)), which is qualitatively consistent with metadynamics simulations (Fig.S10(c)). These results indicate that DiffPIE successfully incorporates protein-gold interactions into protein structure generation.

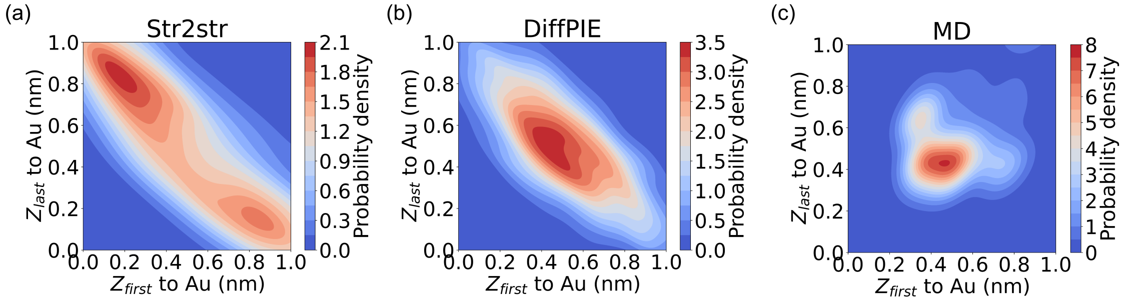


Figure S10: Distance of the first and last residues from the Au(111) surface for structures generated by Str2Str, DiffPIE, and metadynamics simulations. (a) Distributions from Str2Str. (b) Distributions from DiffPIE. (c) Distributions from with metadynamics simulations.

### Details of Side Chain Generations for $A\beta_{16-22}$

We want to emphasize that possible clashes can arise even after minimizing $E_{\mathrm{ab}}$. Minimizing $E_{\mathrm{ab}}$ provides optimal atomic positions of the backbone without involving side chains. For reconstructing side chains of $A\beta_{\text{16-22}}$ on the gold surface, we have tried FASPR to generate initial side chain conformations. However, the default FASPR library does not account for interactions with the Au(111) surface, often resulting in steric clashes and unfavorable side chain orientations at the interface. To address this limitation, we reconstructed side chains from metadynamics simulations onto the DiffPIE-generated backbones, using a strategy analogous to that employed for stapler molecule assembly. Specifically, we aligned the backbone atoms of residue $i$, $\{\mathbf{x}_{\mathrm{C}}^{(i)}, \mathbf{x}_{\mathrm{C}\alpha}^{(i)}, \mathbf{x}_{\mathrm{N}}^{(i)}\}$ to metadynamics samples of dipeptide molecules with the same amino acid type $\{\mathbf{x}_{\mathrm{C}}^{(i),MD}, \mathbf{x}_{\mathrm{C}\alpha}^{(i),MD}, \mathbf{x}_{\mathrm{N}}^{(i),MD}\}$. $\mathbf{x}_{\mathrm{C}}^{(i)}$, $\mathbf{x}_{\mathrm{C}\alpha}^{(i)}$, and $\mathbf{x}_{\mathrm{N}}^{(i)}$ are coordinates of carbonyl carbon, C$\alpha$, and amide $N$ from Str2Str while $\mathbf{x}_{\mathrm{C}}^{(i),MD}$, $\mathbf{x}_{\mathrm{C}\alpha}^{(i),MD}$, and $\mathbf{x}_{\mathrm{N}}^{(i),MD}$ are coordinates of carbonyl carbon, C$\alpha$, and amide $N$ from metadynamics. With the alignment method, only side chain atoms from the MD frames with low backbone RMSD to Str2Str backbone can be selected as candidates. This step significantly reduced steric clashes during side chain replacement.

The matching has been down by minimizing the following loss function:

$$\mathcal{L}^{(i)} = \mathrm{RMSD}(\{\mathbf{x}_{\mathrm{X}}^{(i)}\}, \{\mathbf{x}_{\mathrm{X}}^{(i),MD}\}) + \lambda \|\mathbf{d}^{(i)} - \mathbf{d}^{(i),MD}\|^2. \tag{S.3}$$

$\{\mathbf{x}_{\mathrm{X}}^{(i)}\} = \{\mathbf{x}_{\mathrm{C}}^{(i)}, \mathbf{x}_{\mathrm{C}\alpha}^{(i)}, \mathbf{x}_{\mathrm{N}}^{(i)}\}$ and $\{\mathbf{x}_{\mathrm{X}}^{(i),MD}\} = \{\mathbf{x}_{\mathrm{C}}^{(i),MD}, \mathbf{x}_{\mathrm{C}\alpha}^{(i),MD}, \mathbf{x}_{\mathrm{N}}^{(i),MD}\}$. $\mathbf{d}^{(i)} = (d^{(i-1)}, d^{(i)}, d^{(i+1)})$ where $d^{(i)}$ is the distance between C$\alpha$ of the residue $i$ to the gold surface. $\mathbf{d}^{(i),MD} = (d1, d2, d3)$. $d1$, $d2$, and $d3$ are distances between the gold surface to the methyl carbon of the acetyl cap, the $\alpha$-carbon, and the methyl carbon of the N-methylamide cap in the dipeptide metadynamics simulation. To balance steric clash avoidance with alignment fidelity, we chose $\lambda = 0.5$. This weighting scheme prioritizes the reduction of steric clashes while allowing for moderate deviations in side chain orientation to maintain reasonable structural compatibility. As illustrated in Fig. S11(a), the side chains generated by FASPR frequently clash with the Au surface. Although adjusting the COM distance of the peptide to the Au surface reduces clashes, it cannot control the orientation of the side chain. Our method overcomes this difficulty by replacing side chains with MD-derived conformations. For instance, after alignment, the phenylalanine (PHE) side chain adopts a parallel orientation relative to the Au surface, which is energetically preferred.

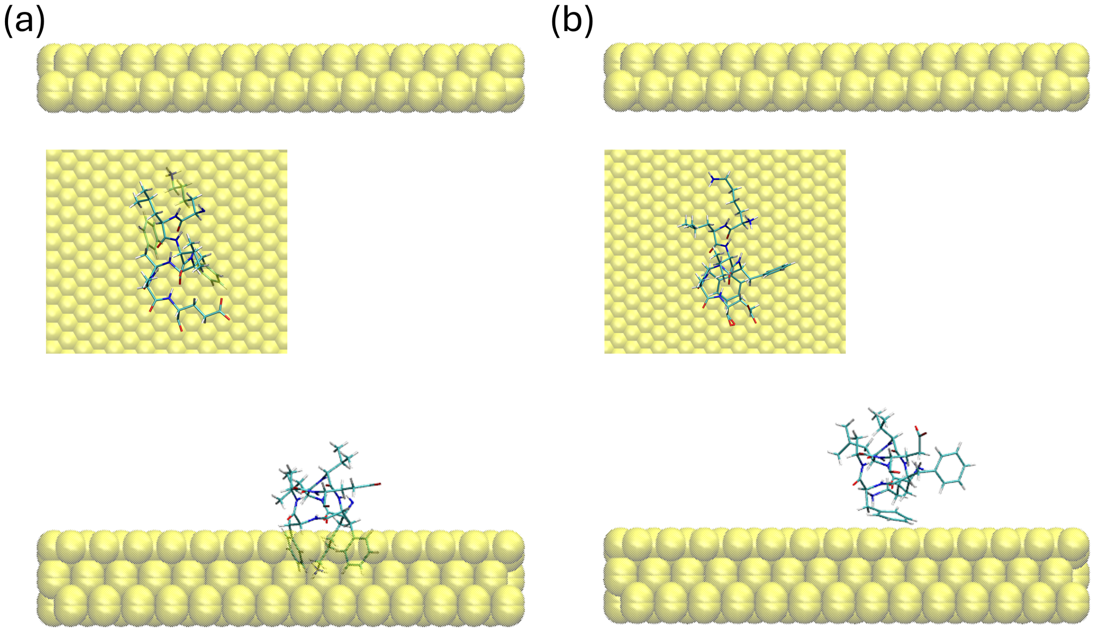


Figure S11: Side chain generation for $A\beta_{16-22}$. (a) A DiffPIE sample with FASPR side chain placed near Au surface with COM-to-Au distance initiated at 0.5 nm. (b) Side chain generated by MD structure replacement after COM-to-Au adjustment. Insets are the top view of the peptide on gold surface.

(4) Iori, F.; Corni, S. Including image charge effects in the molecular dynamics simulations

of molecules on metal surfaces. *Journal of computational chemistry* **2008**, *29*, 1656–1666.
